# Functional importance of JMY expression by Sertoli cells in mediating mouse spermatogenesis

**DOI:** 10.1101/425082

**Authors:** Yue Liu, Jiaying Fan, Yan Yan, Xuening Dang, Ran Zhao, Yimei Xu, Zhide Ding

**Affiliations:** Department of Histology, Embryology, Genetics and Developmental Biology, Shanghai Key Laboratory for Reproductive Medicine, Shanghai Jiao Tong University School of Medicine, Shanghai 200025, China; Center for Experimental Medical Science Education, Shanghai Jiao Tong University School of Medicine, Shanghai 200025, China; Department of Clinical Medicine, Shanghai Jiao Tong University School of Medicine, Shanghai 200025, China

**Keywords:** Sertoli cells, endocytic vesicle trafficking, junction-mediating and regulatory protein (JMY), junction remodeling, blood-testis barrier (BTB), spermatid development

## Abstract

Sertoli cells are crucial for spermatogenesis in the seminiferous epithelium because their actin cytoskeleton supports vesicle transport, cell junction, protein anchoring and spermiation. Here, we show that junction-mediating and regulatory protein (JMY), an actin regulating protein, also affects endocytic vesicle trafficking and Sertoli cell junction remodeling since disruption of these functions induced male subfertility in Sertoli cell-specific *Jmy* knockout mice. Specifically, these mice have: a) impaired BTB integrity and spermatid adhesion in the seminiferous tubules; b) high incidence of sperm structural deformity; c) reduced sperm count and poor sperm motility. Moreover, the cytoskeletal integrity in Sertoli cell-specific *Jmy* knockout mice was compromised along with endocytic vesicular trafficking. These effects impaired junctional protein recycling and reduced Sertoli cell junctions. In addition, JMY interaction with α-actinin1 and Sorbs2 was related to JMY activity and in turn actin cytoskeletal organization. In summary, JMY affects control of spermatogenesis through regulating actin filament organization and endocytic vesicle trafficking in Sertoli cells.

## Introduction

The testicular sustentacular cells, also known as Sertoli cells, are the only somatic cells in the seminiferous epithelium spanning the entire epithelium from the basement membrane into the lumen. They sustain a crucial nursing role through providing physical support, mediating nutrient transport, cleavage and eliciting paracrine signals that affect nascent sperm development (Griswold, 1998; França et al., 2016; Griswold, 2018). These functional roles help account for why sperm production critically depends on functional Sertoli cells interspersed within the seminiferous epithelial layer.

Cell to cell junctions between juxtaposed basal Sertoli cells surround the spermatogonia and delineate the boundary between the apical and basal seminiferous epithelium. These points of contact constituting the blood–testis barrier (BTB) include tight junctions, ectoplasmic specialization (ES), gap junctions and desmosome-like junctions (Mruk and Cheng, 2015). This physical barrier isolates the developing germ cells from immune responses to immune cell infiltration and exogenous toxins trapped in the basal compartment. Therefore, this barrier provides a molecule selectivity filter which creates a unique microenvironment supporting germ cell development (Mital et al., 2011; França et al., 2016; Stanton, 2016). Meanwhile, multiple germ cell–Sertoli cell junctions including desmosome-like junction and the ES, keep germ cells adhering to Sertoli cells. Moreover, the ES localized in the apical region of the Sertoli cells are critical for orienting the elongating spermatid and releasing them once they have developed into mature spermatozoa during spermiation (França et al., 2016; Li et al., 2017). Therefore, the cell junctions elaborated by Sertoli cells, namely both Sertoli-Sertoli cell and Sertoli-germ cell junctions, contribute to sperm production in the seminiferous epithelium.

It is noteworthy that multiple junctions, particularly the tight junctions in Sertoli cells undergo dynamic restructuring during germ cell development. At stage VIII of the seminiferous epithelial cycle during spermatogenesis, preleptotene spermatocytes transit across the BTB near the basement membrane to promote the release of the adherent mature spermatid cells in the apical compartment wherein multiple junctions between adjacent Sertoli cells and Sertoli-germ cells instantaneously break down and then rapidly recombine. It is becoming clear that some tight-, gap-, adherens-, and desmosome-like junctional proteins undergo continuous endocytosis and recycling back to the plasma membrane, thereby resulting in scavenging of the old junctions and replacing them with new junctions (Mruk and Cheng, 2004; Du et al., 2013; Vogl et al., 2014). During this process, regulation of endocytic trafficking of junctional proteins provides a way of rapidly restructuring cell–cell junctions. Failure in any of these steps during endocytosis by Sertoli cells can lead to structural cell junctional defects in the seminiferous epithelium and subsequently an unstable environment for spermatogenesis (Cheng et al., 2011; Jia et al., 2017).

Endocytosis by Sertoli cells is regulated by androgens and cytokines (such as TNF-α, TGF-β2, TGF-β3, IL-6). Androgens can promote endosomal recycling and enhance formation of new cell junctions. In contrast, TGF-β, TNF-α and IL-6 linked signaling promote the fusion of endosomes to lysosomes, which enhances the degradation of cell junctions (Yan et al., 2008; Su et al., 2010). Moreover, the actin cytoskeleton is essential for endocytosis and also for endosomal movement. Numerous studies demonstrated the important roles of both the actin cytoskeleton and actin regulating proteins in trafficking of junctional proteins and maintaining cell junction integrity in Sertoli cells (Li et al., 2016). For instance, recent studies showed that a class of actin-related proteins and their regulators, such as actin-related protein 3 (Arp3) and Wiskott-Aldrich syndrome protein (WASP), were involved in maintaining Sertoli cells tight junctions and BTB function (Lie et al., 2010; Xiao et al., 2014; Mok et al., 2015). More importantly, there are reports indicating that JMY, a junction-mediating and regulatory protein, is a member of the WASP family and it has a similar function to WASP and WHAMM (WASP homolog associated with actin, membranes, and microtubules). In addition, JMY is a multifunctional protein with roles in transcriptional co-activation of p53 and regulation of actin nucleation, which accompanies activation of the Arp2/3 protein complex (Roadcap and Bear, 2009). Analysis of the JMY protein sequence shows that its N-side contains a coiled-coil (CC) domain that can combine with the p300 protein. On the other hand, the C-side has three WASp Homology 2 (WH2) domains. They can combine with actin and a connector and an acidic (CA) domain and in turn with the Arp2/3 complex. When in the cytosol, JMY can activate Arp2/3 and produce branched F-actin filaments along with unbranched filaments via its tandem WH2 domains (Coutts et al., 2009; Zuchero et al., 2009; Rottner et al., 2010).

It is uncertain whether JMY has a functional role in mediating spermatogenesis because its expression pattern has not been described in Sertoli cells. In addition, the effects are unknown of JMY protein downregulation or loss of function in Sertoli cells. Herein, we report that the JMY protein is highly expressed in Sertoli cells and is an actin regulator. This function is crucial in the maintenance of cell junction integrity which is vital for preserving spermatids adhesion and BTB integrity in Sertoli cells.

## Results

### Expression and localization of JMY in Sertoli cells

Immunofluorescent staining and immunoblotting detected JMY expression in the seminiferous epithelium of mouse testis and primary cultured Sertoli cells (Fig. 1 A, Fig. 2A, Fig. S1). JMY expression was prominent in Sertoli cells whereas it was much less evident in spermatids throughout the seminiferous epithelium (Fig. 1 A, Fig. S1). These results are consistent with Western blot analysis which also identified JMY expression in the testis and Sertoli cells (Fig. 2 A). Additionally, JMY was localized in the cytosol of cultured Sertoli cells and it was somewhat more evident at the F-actin-rich domain (Fig. 1 B), which is consistent with its actin-regulatory involvement.

**Figure 1.**
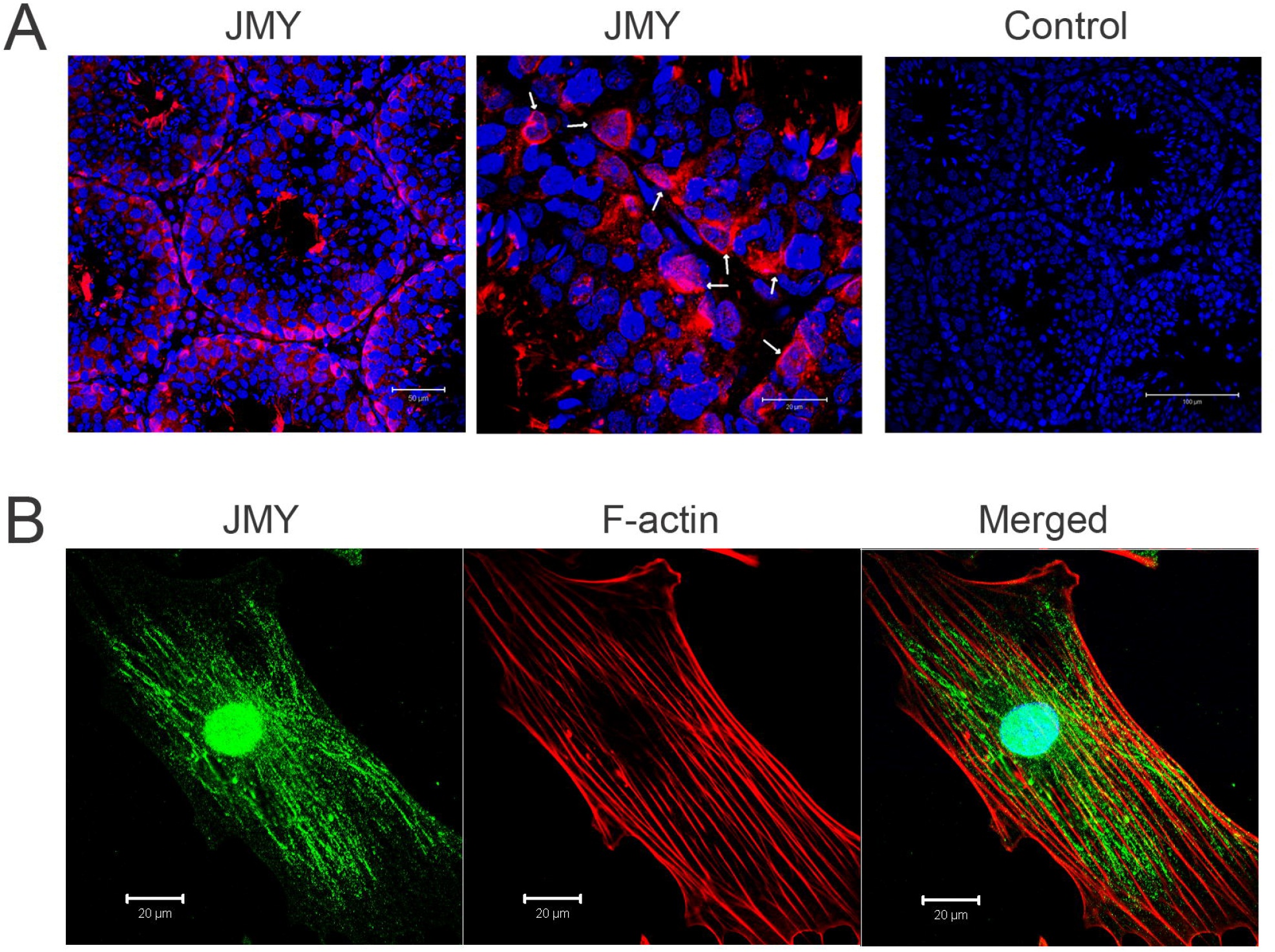
JMY expression in testis and Sertoli cells. (A) JMY Immunofluorescent staining shows that JMY expression is localized in the Sertoli cells of mouse testis (Arrows indicated). Bars: (main) 50 μm; (enlarged) 20 μm. (B) JMY immunofluorescent staining and F-actin rhodamine-phalloidin staining of primary cultured Sertoli cells shows that its expression is localized in the cytosol and nucleus. In the cytosol, JMY has a filamentous distribution, which in some places colocalizes with F-actin. Scale Bar: 20μm.

**Figure 2.**
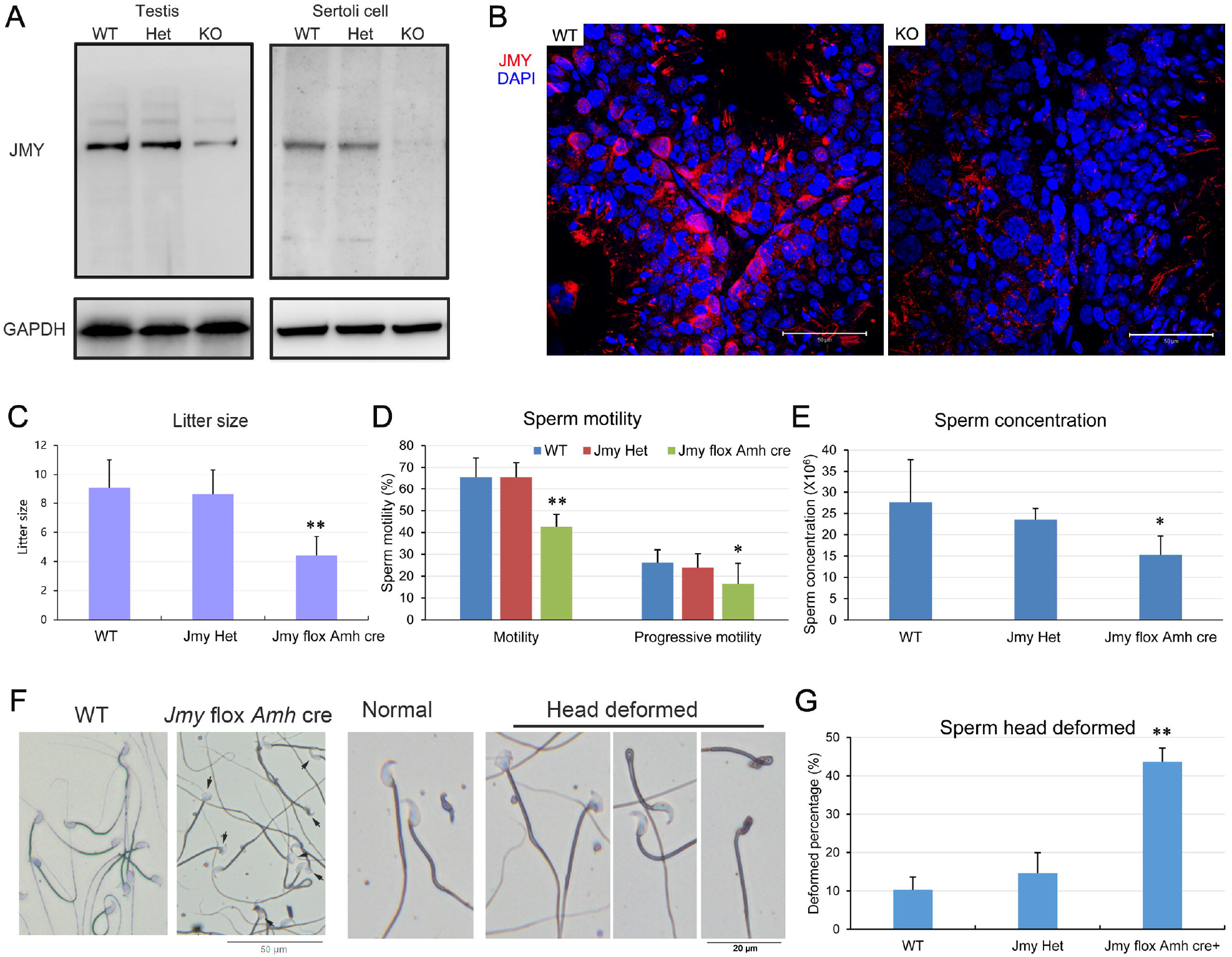
WT and *Jmy* CKO sperm morphology analyses. (A) Western blot analysis resolves JMY expression in mouse testicular Sertoli cells. JMY expression was greatly reduced in CKO testis relative to that in its WT counterpart and Het testis, JMY expression was undetectable in Sertoli cells of CKO mice. GAPDH expression validated loading equivalence. (B) JMY immunofluorescent staining of shows that JMY expression in Sertoli cells was absent in testis from *Jmy* CKO mice. Scale Bar: 50 μm. (C) Litter size of mature female mice mated with *Jmy* CKO male mice was greatly reduced relative to those mated with WT and Het male mice. Error bars represent SD (n=10). \*\**P* <0.01. (D) Sperm motility analyzed by CASA shows that both sperm motility and progressive motility of *Jmy* CKO mice were less than that in WT and Het mice. Error bars represent SD (n=5). \**P* < 0.05; \*\**P* <0.01. (E) Sperm concentration analyzed by CASA, shows that sperm concentration of *Jmy* CKO mice was lower than that in WT and Het mice. Error bars represent SD (n=5). \**P* < 0.05. (F) Sperm morphology of WT and *Jmy* CKO mice. Except for the prominent deformations in the *Jmy* CKO sperm heads, the other regions appear unchanged from the normal WT morphology (CKO, arrows indicated). Bars: (main) 50 μm; (enlarged) 20 μm. (G) Deformed sperm head ratios calculated from five independent experiments of at least 200 spermatozoa in each independent experiment. Error bars represent SD (n=5). \*\**P* <0.01.

### Conditional knock out (CKO) of *Jmy* in the Sertoli cells

To identify the function of JMY in Sertoli cells, *Jmy* loxP mice were constructed. Accordingly, two loxP sites were inserted into the flanking regions of the third exon in the *Jmy* gene using homologous recombination (Fig. S2). Then, the *Jmy* flox (*Jmy*^loxP/loxP^) mice were mated with *Amh* cre mice in order to generate the expected Sertoli cell specific *Jmy* CKO mice (*Jmy* ^loxP/loxP^; *Amh* cre^+^). They carried a Sertoli cell specific *Jmy* deletion which was obtained by splicing out exon 3 using the Sertoli cell specific expression of Cre recombinase. In these *Jmy* CKO mice, Western blot and immunofluorescent analysis documented that JMY testicular expression was absent in Sertoli cells (Fig. 2 A and B), which confirms *Jmy* gene knockout in these mice.

### Reduced male fertility and poor sperm quality in *Jmy* CKO mice

To assess the fertility of the *Jmy* CKO mice, each male was mated with two mature females for one week and then litter sizes were compared with their wild type (WT) counterpart. As expected, the litter size of male *Jmy* CKO mice (4.65 ± 1.31) was significantly less than that of both the WT and the heterozygote (Het, *Jmy* lox/-) males (WT: 7.40 ± 2.01, Het: 7.29 ± 0.95; Fig. 2 C). Moreover, the *Jmy* CKO sperm motility and concentration also decreased in comparison with those of the WT and the Het males (Sperm motility in WT: 65.60 ± 8.76, Het: 66.20 ± 6.58, CKO: 42.50 ± 5.74. Sperm progressive motility in WT: 26.20 ± 5.89, Het: 23.80 ± 6.54, CKO: 16.50 ± 6.14. Sperm concentration, ×10^6^ per ml, in WT: 27.62 ± 7.82, Het: 23.54 ± 2.62, CKO: 15.28 ± 4.41. Fig. 2 D and E), while the sperm head deformed ratios were greatly enlarged (WT: 10.23 ± 3.38, Het: 14.60 ± 5.33, CKO: 43.65 ± 3.55. Fig. 2 F and G). Thus, the reduced litter size and poor sperm quality clearly indicate that male subfertility was established in the *Jmy* CKO mice.

### Morphological alteration in *Jmy* CKO testes

The testicular morphology of *Jmy* CKO mice was abnormal compared to that in the WT and the Het counterparts. The *Jmy* CKO testes had asyntactic Sertoli cells at the basal region of the seminiferous epithelium and poor adhesion between spermatids and Sertoli cells, leading to the abscission of round spermatids into the lumen of seminiferous tubules (Fig. 3 A), whereas in the sections of *Jmy* CKO caput epididymis, masses of round spermatids were evident in the epididymal duct lumen (Fig. 3 B).

**Figure 3.**
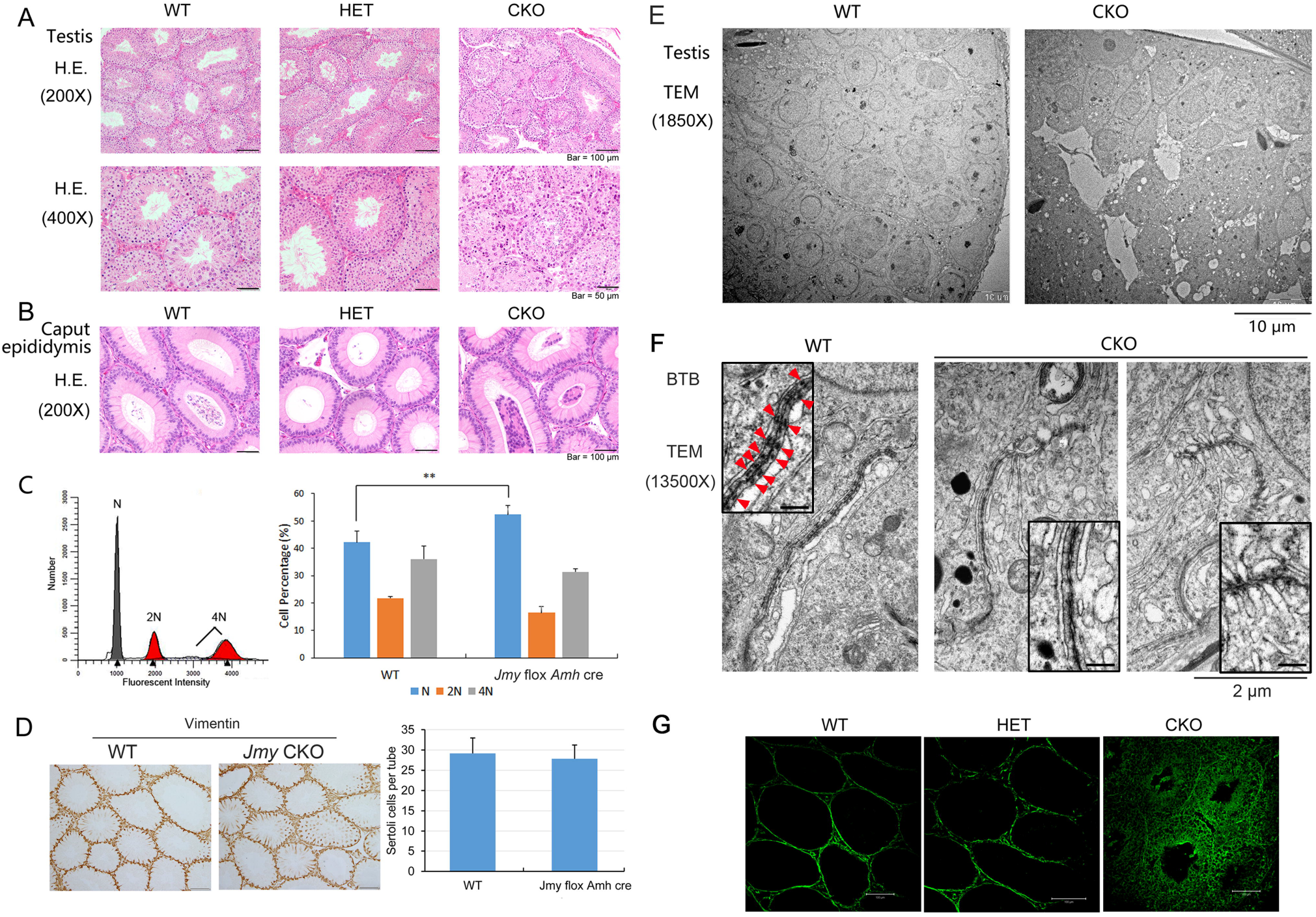
Loss of JMY function induces testicular histological disorder. (A-B) In H&E stained testicular (A) and caput epididymal (B) sections, testicular cellular disorganization is evident and spermatid epithelial adherence is less. These alterations obstruct caput epididymis lumen patency in *Jmy* CKO mice. Bars: (main) 100 μm; (enlarged) 50 μm. (C) Cell cycle analysis results resolve different cell types in the seminiferous epithelium. DNA content of indicated cell types: diploid (2N), tetraploid (4N) and haploid (N). Statistical results show that the proportion of haploid cells increased in *Jmy* CKO mice. Error bars represent SD (n = 5; 30,000 cells per test). \*\**P*<0.01. (D) Vimentin immunohistochemical staining is evident in testicular sections. Vimentin is one kind of intermediate filamentous protein that is an established marker of Sertoli cells. Sertoli cell numbers in every seminiferous tubule from WT mice and *Jmy* CKO mice are not significantly different from one another. Error bars represent SD (n = 30; 5 tubules/trial). (E-F) Transmission electron micrograph analysis of seminiferous epithelium shows that spermatid cell adhesions are incomplete within the seminiferous epithelium of *Jmy* CKO testes (E) which are accompanied by disrupted junctional structures between neighboring Sertoli cells (F). Arrows in (F) indicate the actin bundles of the ES in WT mice. Bars: (E) 10 μm; (F) 2μm; (zoom in F) 0.2 μm. (G) The biotin assay evaluated BTB integrity in vivo. Alexa Fluor 488 staining visualized biotin distribution. The diffuse green fluorescent staining throughout all the layers of the testicular seminiferous epithelium indicates that loss of JMY function contributes to defective BTB structural formation in the he *Jmy* CKO mice. Scale Bar: 100 μm.

To characterize the cell type alterations in the seminiferous epithelium, cell cycle analysis was performed in *Jmy* CKO and WT mice. The cell types were classified according to karyotype analysis based on comparing spermatogonia, secondary spermatocyte and Sertoli cells of diploid cells with those of: a) primary tetraploid spermatocytes in the tetraploid cells; b) haploid spermatids and spermatozoa. The results revealed a significantly higher percentage of haploid cells in *Jmy* CKO seminiferous epithelium compared to that in the WT seminiferous epithelium (Fig. 3 C). Meanwhile, the number of Sertoli cells was evaluated based on the immunostaining expression pattern of vimentin, which is a Sertoli cell marker, in 8 weeks old *Jmy* CKO and WT mice. The number was essentially the same between these two groups (Fig. 3 D).

Additionally, electron micrographs showed that cell adhesions were unconsolidated within the seminiferous epithelia of the *Jmy* CKO testes (Fig. 3 E), and junctional structures between the Sertoli cells, especially the actin bundles of the ES were greatly disrupted (Fig. 3 F).

The in vivo classical biotin permeation assay compared the BTB integrity in the *Jmy* CKO with that in the WT and Het testes. As expected, the staining patterns clearly defined basal spermatogonia but none were present in the upper apical cell layers in the WT and Het testes. In contrast, biotin readily permeated throughout the entire germinal epithelium in the *Jmy* CKO testes (Fig. 3 G).

In summary, loss of *Jmy* in Sertoli cells impaired Sertoli cell junctional barrier formation and loosened spermatid attachment.

### Disrupted cell junctions and endocytosis in *Jmy* CKO testes

To confirm the alteration of cell junctions in *Jmy* CKO testes, immunoblot analyses evaluated junctional signature protein expression. ZO-1, claudin11 (tight junction proteins), as well as β-catenin (adherens junction protein) expression levels decreased in the *Jmy* CKO testes in comparison to those in WT testes (Fig. 4 A). Meanwhile, immunohistochemistry analyses showed that expression of N-cadherin, specifically localized in the basolateral region of adjacent Sertoli cells in WT testes decreased and was more diffuse in the junctional compartment of Sertoli cells in *Jmy* CKO testes (Fig. 4 B). Thus, these results show that the declines in expression of cell junctional proteins impaired tight junctioncontacts and BTB integrity between neighboring Sertoli cells in the *Jmy* CKO testes.

**Figure 4.**
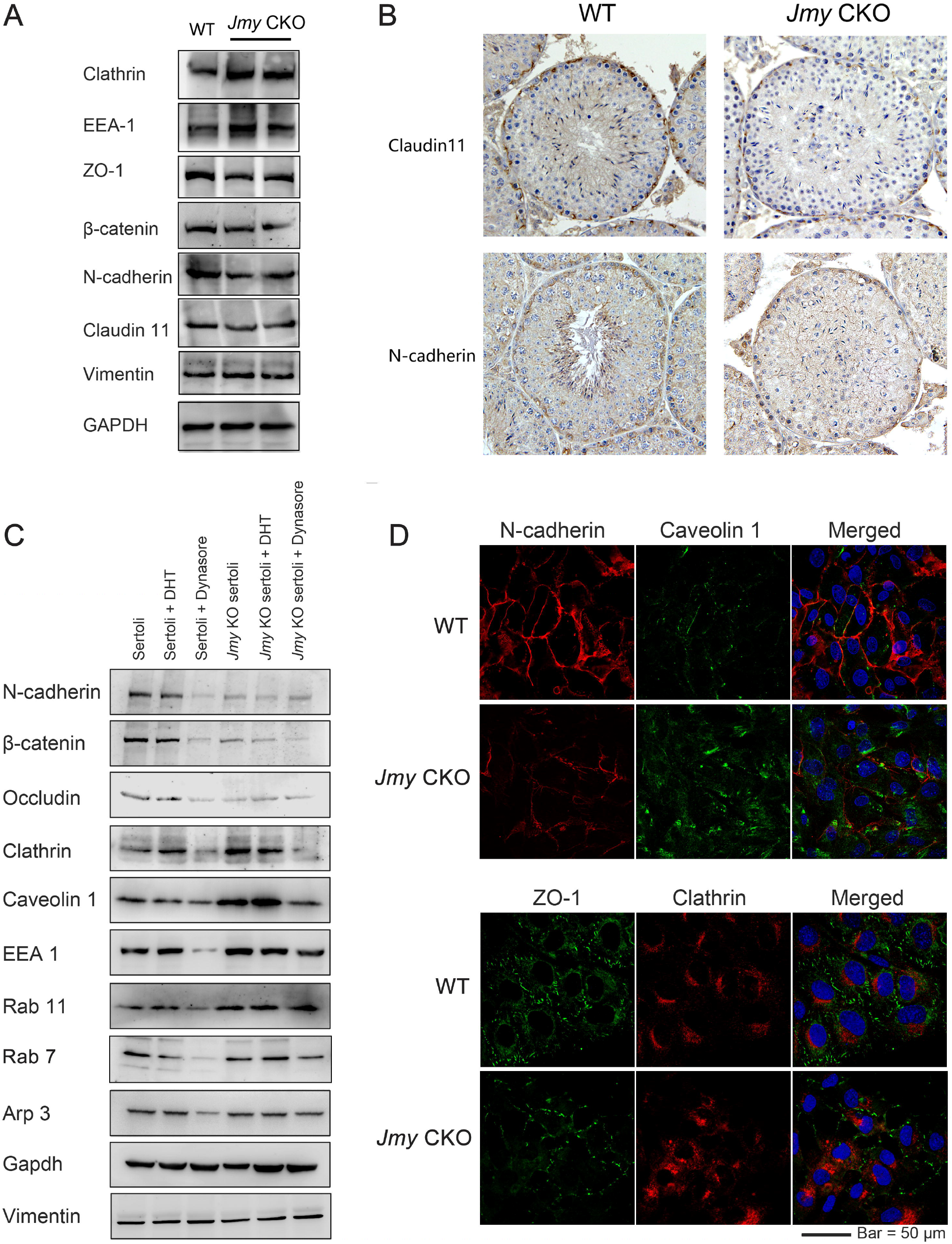
Loss of JMY function disrupts cell junctional integrity between testicular Sertoli cells. (A) Comparison of Western blot analyses of testicular protein expression in WT and *Jmy* CKO mice. Clathrin and EEA-1, endocytosis related proteins; ZO-1, β-catenin, N-cadherin and Claudin11 expressions identified junctional proteins; Vimentin and GAPDH expression validated loading control equivalence. (B) Immunohistochemical staining of testicular sections, junctional Claudin11 and N-cadherin proteins, shows that their expression is localized to the basolateral region of adjacent Sertoli cells in WT testes. In contrast, such delimited junctional expression is not evident in *Jmy* CKO testicular Sertoli cells. Scale Bar: 50 μm. (C) Western blot analyses of proteins expression patterns in WT Sertoli cells and *Jmy* CKO Sertoli cells, as well as in those treated with either DHT or Dynasore. Vimentin and GAPDH expression validated loading control equivalence. (D) Immunofluorescent staining of junction proteins (N-cadherin and ZO-1) and endocytosis related proteins (Caveolin-1 and Clathrin), shows decreased amount of N-cadherin and ZO-1 expression whereas caveolin-1 and clathrin expression increased in the *Jmy* CKO Sertoli cells. Scale Bar: 50 μm.

Notably, dynamic trafficking of tight junctionproteins is crucial for maintaining barrier function and it is greatly dependent on endocytosis. Endocytic trafficking of junctional proteins, including continuous endocytosis and recycling back to the plasma membrane enables rapid junctional remodeling. Immunoblot analysis of clathrin (an endocytic vesicle marker) and EEA-1 patterns (an early endosomal marker) showed that they increased in *Jmy* CKO testes (Fig. 4 A). These changes indicate that decreased levels of junctional protein expression may result from endocytosis up-regulation in *Jmy* CKO testes.

### Disrupted cell junctions and endocytosis in *Jmy* CKO Sertoli cells

Primary cultured Sertoli cells were used to confirm that alterations in cell junctional integrity and endocytosis occur in *Jmy* CKO mice. As stated above, immunoblot analyses showed that the tight junction protein (occludin) and the adherens junction proteins (β-catenin and N-cadherin) were downregulated in *Jmy* CKO Sertoli cells. In contrast, the endocytic vesicle markers (clathrin and caveolin 1), the early endosomal marker (EEA-1) and the recycling endosome marker (Rab 11) were upregulated in *Jmy* CKO Sertoli cells, whereas the later endosome marker (Rab 7) was unchanged in *Jmy* CKO Sertoli cells (Fig. 4 C). Immunofluorescent staining analyses of cultured Sertoli cells also confirmed declines in N-cadherin and ZO-1 localization at the cell–cell interface of *Jmy* CKO Sertoli cells, as well as the increases in contents of caveolin 1 and clathrin in *Jmy* CKO Sertoli cells (Fig. 4 D). These findings unequivocally confirm losses in cell junctional formation whereas endocytosis increased in *Jmy* CKO Sertoli cells. These changes suggest that JMY expression contributes to controlling endocytic trafficking of junctional proteins.

Another indication of JMY involvement is that treatment of normal WT Sertoli cells with dihydrotestosterone (DHT), markedly upregulated clathrin, an activator of endocytosis and endosome recycling, and EEA-1 and Rab11 endocytosis related proteins. These changes are consistent aforementioned indications of JMY involvement in controlling BTB integrity and endocytic trafficking. Likewise, upon treatment of WT Sertoli cells with dynasore, an inhibitor of dynamin and vesicle trafficking, the contents of junctional proteins, i.e. occludin, β-catenin and N-cadherin significantly decreased, which is also consistent with their alterations in *Jmy* CKO Sertoli cells (Fig. 4 C). Therefore, the effects of loss of JMY function on Sertoli cells are similar to those induced by DHT and dynasore treatments. This correspondence strengthens the notion that JMY is involved in endocytic vesicle trafficking and endosomal recycling.

### Disrupted endosomal cycling in *Jmy* CKO Sertoli cells

Endocytosis occurs through multiple steps which initially include internalization of proteins spanning the entire width of the plasma membrane followed by vesicular trafficking into early endosomes. Subsequently, the endosomes are inserted into the lysosomal degradative pathway or recycled back to the plasma membrane. To identify the trafficking fate of the endocytosed junctional proteins, we used a biotinylation assay that labels extracellular amino residues of membrane proteins and allows their trafficking to be monitored. In WT Sertoli cells, the biotin labeled proteins were initially detected at the surface of cells and endocytosis gradually moved them into the cytoplasm. Because DHT induces endosomal recycling, the biotin labeled proteins recycled back to the plasma membrane of Sertoli cells at the cell-cell interface after either 1 hour or 3 hours culture in vitro (Fig. 5 A). Even though biotin-labeled membranous proteins were transported into the cytoplasm by endocytosis in *Jmy* CKO Sertoli cells, endocytic trafficking was not completed since the proteins accumulated in the cytoplasm (Fig. 5 A). Alternatively, treating with dynasore blocked endocytosis at a step prior to biotin labeling in *Jmy* CKO Sertoli cells, thus the biotin labeled membranous proteins remained on the cell surface (Fig. 5 B).

**Figure 5.**
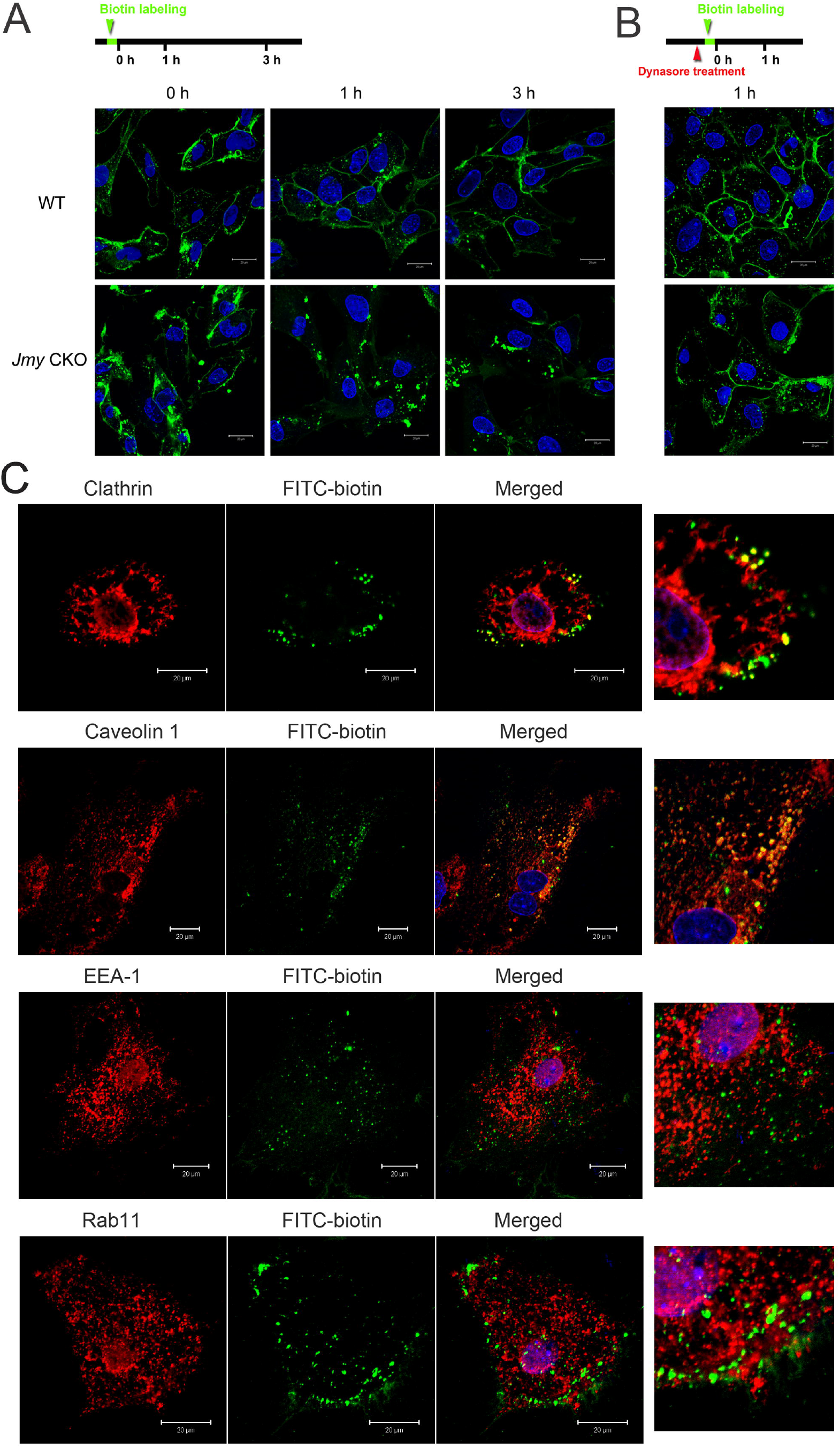
Endosome cycling disrupted in *Jmy* CKO Sertoli cells. (A) Changes in cell surface biotinylation evaluated endocytosis and recycling in Sertoli cells. After biotin labeling for 0h, 1h and 3h, the intracellular translocation of the biotinylated membrane proteins was detected by fluorescent staining. Bars: 20 μm. (B) Dynasore treatment blocked endocytosis, which resulted in less cytoplasmic biotinylated protein accumulation in the cytoplasm. Bars: 20 μm. (C) The contribution by functional JMY expression to mediating endocytosis of biotinylated membrane proteins was evaluated based on its co-localization with endosomal biomarkers. In Jmy CKO Sertoli cells, immunofluorescent staining shows that the biotinylated membrane proteins accumulated in the cytoplasm and co-localized with endocytic vesicle markers (clathrin and caveolin), whereas less biotinylated proteins co-localized with endosomal protein markers (EEA-1 and Rab 11). Scale Bars: 20 μm.

Moreover, immunofluorescent staining also demonstrated that a large amount of biotin labeled proteins accumulated in the cytoplasm of *Jmy* CKO Sertoli cells and colocalized with endocytic vesicle markers, clathrin and caveolin 1, rather than the endosomal markers, EEA-1 and Rab 11 (Fig. 5 C). These results suggest that the loss of *Jmy* function blocks vesicle trafficking into the early endosomes.

### Identification of proteins interacting with JMY by immunoprecipitation (IP) and liquid chromatography tandem mass spectrometry (LC-MS) analysis

Due to the role of JMY as an actin regulator, we hypothesized that cooperative interaction between JMY and some other proteins contribute to endocytic vesicle trafficking. To gain insight into the molecular mechanism underlying JMY effects on vesicle trafficking, the proteins interacting with JMY in Sertoli cells were separated by IP and then identified by LC-MS analysis (Table S1). Kyoto Encyclopedia of Genes and Genomes (KEGG) databases (http://www.genome.jp/kegg/) was used to analyze the MS data in order to search for functional annotation terms and pathways. The analysis showed that the proteins interacting with JMY are mainly constituents of a group classified as “Regulation of actin cytoskeleton”, “Tight junction”, “Focal adhesion” and “Endocytosis” in Sertoli cells (Table S2). Similarly, Clusters of Orthologous Groups of proteins (COG) analysis revealed that 15 proteins identified by MS were classified as “Cytoskeleton” (Fig. S3). Considering the known actin-regulating activity of JMY, such we were prompted to determine if it interacts with these other actin cytoskeleton related proteins. Within this group, drebrin, actin-related protein 2/3 complex (Arp2/3), Anxa2, Cofilin, Gelsolin, spectrin have been reported to participate in regulating cell junction formation and function in Sertoli cells (Guttman et al., 2002; Lui et al., 2003; Lie et al., 2010; Li et al., 2011; Aristaeus de Asis et al., 2013; Chen et al., 2017; Chojnacka et al., 2017).

Previous studies demonstrated that the WH and CA domains of JMY bound to the actin and Arp2/3 complex to promote actin nucleation (Coutts et al., 2009; Zuchero et al., 2009; Rottner et al., 2010). Herein, our analysis predicted that JMY interacts with the Arp2/3 complex as well as other actin binding protein candidates that include α-actinin1, Sorbin and SH3 domain containing protein 2 (Sorbs2) in Sertoli cells. If this prediction proves to be valid, the structure of the actin and Arp2/3 complex may in fact also contain other constituents.

### Verification of interactions between JMY and two actin binding proteins, α-actinin1 and Sorbs2

Since α-Actinin1 and Sorbs2 very prominently interact with JMY based on their presence in immunoprecipitates of JMY, we presumed that they support JMY function by serving as its co-factors. To confirm such an interaction, we interrogated immunoblots obtained with JMY antibodies for both α-actinin1 and Sorbs2. Their presence in the JMY immunoprecipitates shown in Figure 6A documents that JMY interacts with α-actinin1 and Sorbs2.

**Figure 6.**
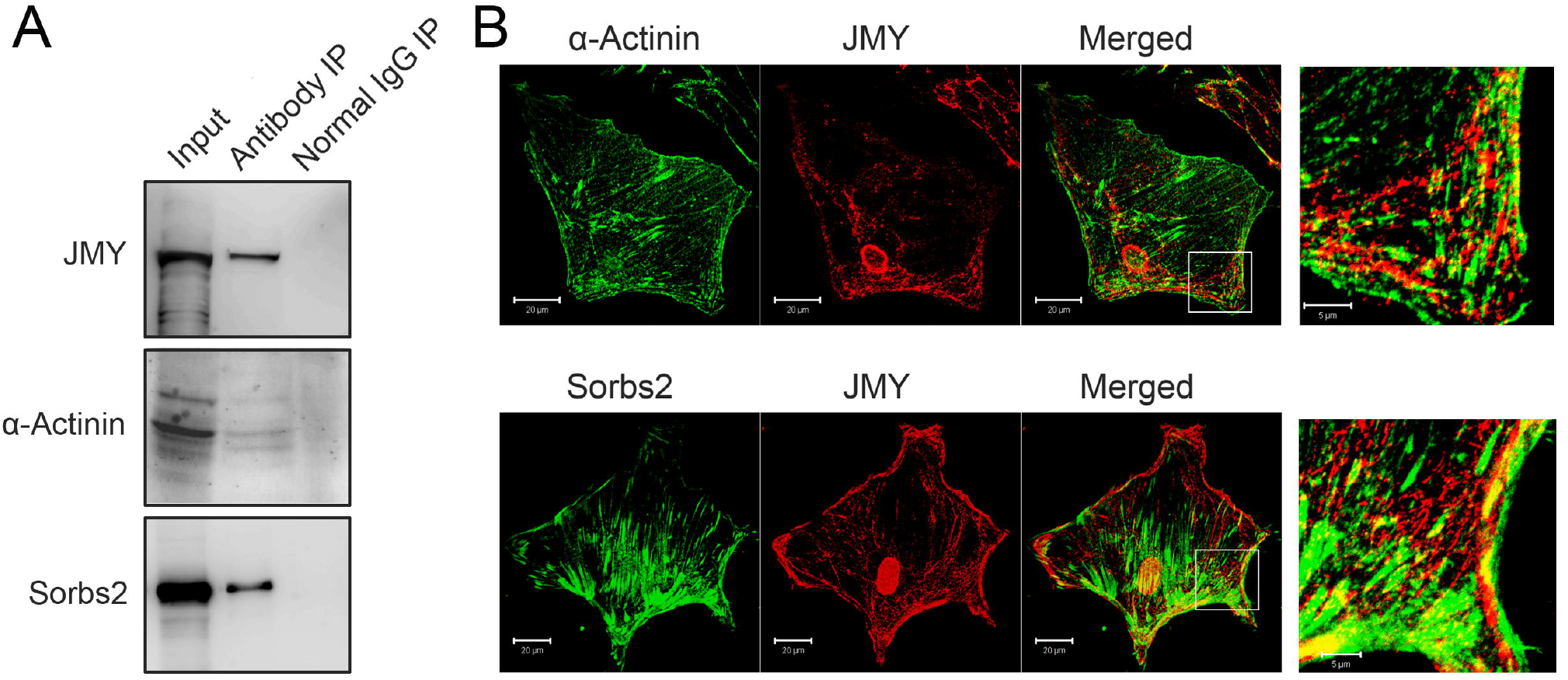
Identification of the proteins interacting with JMY in Sertoli cells. (A) Proteins coimmunoprecipating with JMY antibodies are evident in Western blots interrogated with α-actinin1 and Sorbs2 antibodies in a protein lysate of Sertoli cells expressing JMY. (B) Immunofluorescent staining identifies co-localization of JMY with either α-actinin1 or Sorbs2 in Sertoli cells. Scale Bars: (main) 20 μm; (zoom) 5 μm

Furthermore, immunofluorescent staining substantiates α-actinin1 and Sorbs2 colocalization in cultured Sertoli cells. The images clearly show that they are present in close proximity to the actin cytoskeleton. More importantly, JMY colocalizes at the cell edges with this complex formed between α-actinin1 and Sorbs2 and the actin cytoskeleton network (Fig. 6 B). Colocalization of JMY with these proteins strengthens the notion that these proteins bind to JMY. Accordingly, it is possible that they may interact with one another and mediate specific biological functions such as endocytosis.

### Differential expression of α-actinin 1 and Sorbs2 in *Jmy* CKO Sertoli cells

JMY involvement in fostering a structured actin network was confirmed by the fact that rhodamine-phalloidin stained actin filaments were disordered with highly branched and extensive foci in *Jmy* CKO Sertoli cells (Fig. 7 A). Nevertheless, the expression and localization of Arp3 in *Jmy* CKO Sertoli cells were clearly unchanged from those in WT Sertoli cells (Fig. 4 C, Fig. 7 B and C). This invariance suggests that the disorganized and unstructured actin cytoskeleton induced by loss of *JMY* function was not dependent on an interaction between JMY and Arp2/3.

**Figure 7.**
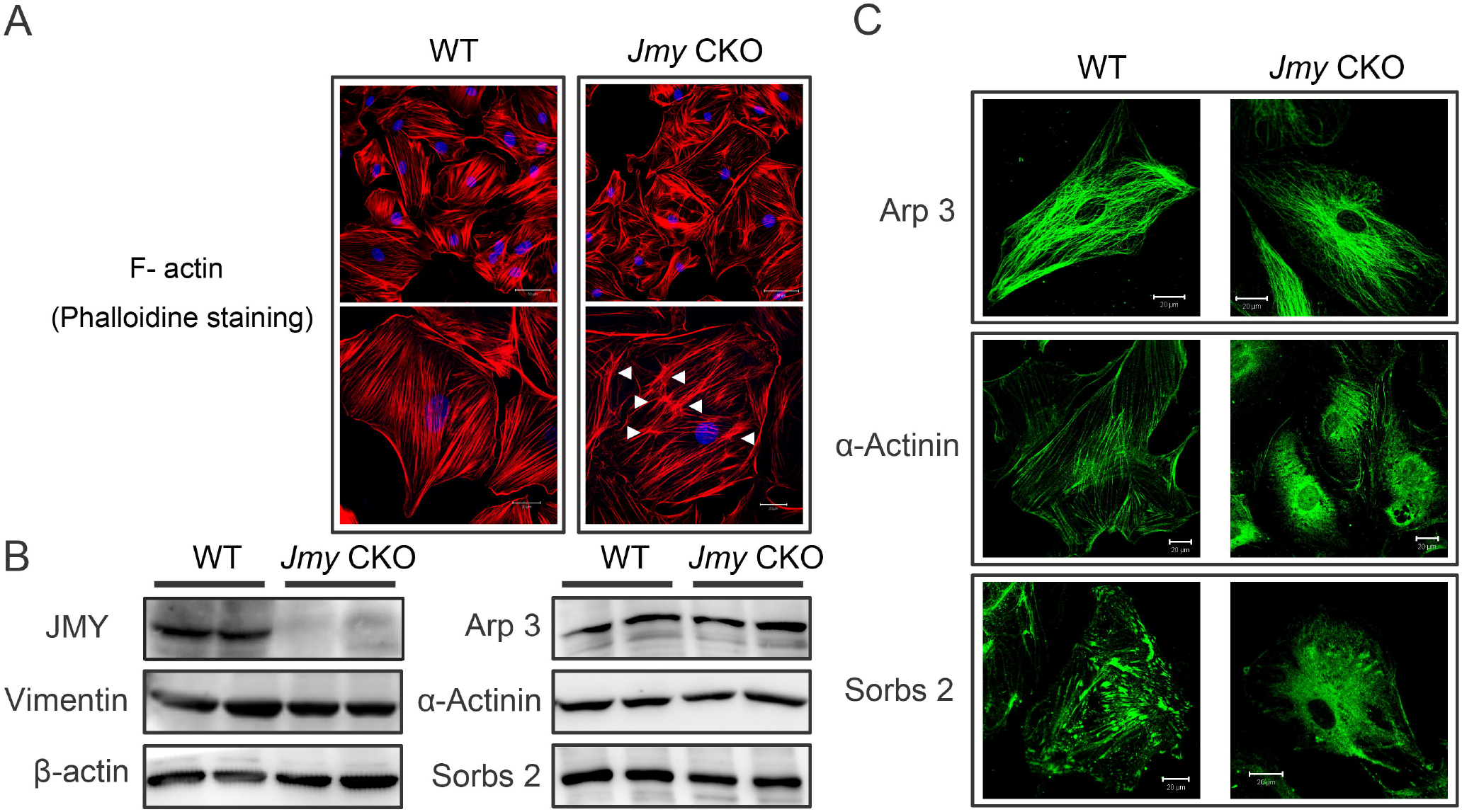
Loss of JMY function disrupts interactions between actin filament and actin regulating proteins in Sertoli cells. (A) Rhodamine-phalloidin staining identified numerous actin filaments foci in *Jmy* CKO Sertoli cells (Arrows highlight). Scale Bars: (main) 50 μm; (enlarged) 20 μm. (B) Western blot analyses of α-Actinin and Sorbs2 expression show that their expression levels are slightly less in *Jmy* CKO Sertoli cells than in the WT counterpart, but that of Arp3 is unchanged in *Jmy* CKO Sertoli cells. Vimentin and ß-actin expression levels document loading control equivalence. (C) Localized Arp3, actinin1 and Sorbs2 immunofluorescent staining in WT Sertoli cells and *Jmy* CKO Sertoli cells. Arp3, actinin1 and Sorbs2 colocalize with actin filaments in WT Sertoli cells whereas their close proximity to one another is lost in *Jmy* CKO Sertoli cells. Scale Bars: 20 μm.

Although both α-actinin1 and Sorbs2 strongly interact with the actin cytoskeleton, their relationship to JMY function required clarification. Notably, in *Jmy* CKO Sertoli cells, both the α-actinin and Sorbs2 expression levels detected by immunoblotting were slightly reduced (Fig. 7 B). A more striking change was that loss of JMY function caused α-actinin 1 and Sorbs2 to lose their close proximity to one another. This disruption is evident based on the altered immunofluorescent staining pattern showing disorganization of the actin cytoskeletal filament network (Fig. 7 A). What is even more informative is that changes in their immunofluorescent staining pattern was delocalized and accompanied disordering of the actin filament organization (Fig. 7 C). Taken together, it is apparent that JMY expression is essential for positioning α-actinin1 and Sorbs2 in locations that are commensurate for mediating development of an organized actin cytoskeleton network.

## Discussion

There is suggestive evidence that the actin cytoskeleton and its binding proteins play an important role in maintaining Sertoli cell function and supporting BTB integrity and spermiogenesis. Here we identify a novel role for JMY, which is an actin regulating protein. It is an essential factor for enabling dynamic remodeling of the junctional proteins mediated by endocytic trafficking in Sertoli cells. JMY involvement in this process is evident because loss of *Jmy* function impaired BTB structural and integrity formation in Sertoli cells, which is obligatory for preventing premature sloughing of developing germ cells from the seminiferous epithelium. Such protection is necessary because it prevents dysfunctional sperm development from reducing sperm fertility as a consequence of declines in sperm count and motility that is associated with a high incidence of sperm head structural deformity.

Sertoli cells are rich in actin filaments, which play a crucial role in certain biological activities, e.g., vesicle transport, ES formation and spermatid head shaping (Du et al., 2013; Vogl et al., 2014; Li et al., 2016; Li et al., 2017). Our results indicate that the JMY distribution pattern is reflective of a possible association between changes in JMY expression and dynamic modulation of the actin cytoskeleton. Specifically, JMY expression was evident in the Sertoli cells and later-stage spermatogenic cells in the seminiferous epithelium. JMY expression exhibited a filament-like expression localized parallel to the actin filaments in Sertoli cells, whereas in the later-stage spermatogenic cells, it was specifically localized in the perinuclear ring and manchette structure which was also rich in actin filaments. Such observations suggested that JMY is likely involved in mediating actin-regulating activity during spermatogenesis. Accordingly, we undertook a characterization of the underlying molecular mechanisms mediating JMY function in spermatogenesis.

To determine the contribution by JMY expression in either Sertoli cells or spermatogenic cells to spermatogenesis, we engineered mice both with Sertoli cell-specific deletion of *Jmy* (*Jmy* flox; *Amh* cre) and germ cell-specific deletion of *Jmy* (*Jmy* flox; *Mvh* cre). Even though the mice with germ cell-specific deletion of *Jmy* were healthy and fertile (Fig. S4), male mice with Sertoli cell-specific deletion of *Jmy* were sub-fertile. This condition was attributable to low sperm count and poor sperm motility as well as appreciable morphological deformity. Disruption of BTB integrity accompanying loss of *Jmy* function could account for declines in sperm fertility since reduced barrier function led to premature sloughing of developing spermatids.

In Sertoli cells, junctional structures undergo extensive organization and transition between “close” and “open” configurations to facilitate transport of germ cells across the BTB as well as control spermatid adhesion or sperm release. This dynamic remodeling of cell junctional barrier function greatly depends on endocytosis and linked endocytic trafficking pathways, which select the endocytosed proteins to be sorted for degradation or recycling (Cheng et al., 2011; Vogl et al., 2014). Moreover, ES is a unique actin-based cell-cell anchoring junction constituted by bundles of actin filaments that are situated between the cisternae of the endoplasmic reticulum and the apposing plasma membrane. In general, these structures are restricted to the interface between Sertoli cells and spermatids and are designated as the apical ES. On the other hand, those located at the Sertoli-Sertoli cell interface are referred to as the basal ES, which usually coexists with tight junctions and gap junctions to constitute the BTB structure. Interestingly, germ cell transport as well as endocytic vesicle trafficking in Sertoli cells requires rapid reorganization of these microfilament bundles, so that they are efficiently converted from a “bundled” to “unbundled/branched” configuration (Li et al., 2015). Briefly, there is no doubt that irrespective of the type of actin bundle configuration, in all cases they support and maintain cell junction integrity. On the other hand, actin branching is a critical step weakening cell junction cohesiveness resulting in junctional reorganization in the seminiferous epithelium. Our electron microscopy analysis confirmed that these alterations occurred in the *Jmy* CKO seminiferous epithelium. Notably, the junctional BTB structures appeared disfigured and actin filament bundle density declined at the basal ES. Such changes cause the BTB to become more permeant resembling those in more leaky epithelia and endothelia. Furthermore, the phalloidin stained actin filaments of *Jmy* CKO Sertoli cells were in disarray having highly branched and enlarged as well as more numerous foci, suggesting that JMY might be essential as well for the maintenance of the effective bundled actin filaments and the ES junctions in Sertoli cells. These changes may also explain why loss of JMY function disrupts cell junction formation; namely, it occurs as a consequence of actin branching.

Additionally, we found that loss of *Jmy* function in Sertoli cells reduced expression of junctional proteins (i.e. ZO-1, claudin11, N-cadherin, occludin and β-catenin) and increased expression of endocytosis related proteins (i.e. clathrin, caveolin 1, EEA-1 and Rab 11) both in the testis and in the Sertoli cells. These effects are tentatively supportive of a mechanism, which links impaired cell junctions and BTB formation with abnormal endocytosis in Sertoli cells. To evaluate this possibility, the effects of testosterone (DHT) treatment or dynasore treatment were determined in an in vitro Sertoli cell culture system since these agents can increase or decrease BTB resistance, respectively and thereby mimic their dynamic behavior in vivo. Our usage of these modulators is based on reports in which testosterone promoted cell junction integrity, apparently via enhancing protein endocytosis and recycling, while dynasore, a well-characterized dynamin inhibitor effectively blocks endocytosis and vesicle trafficking. In our case, testosterone treatment of WT Sertoli cells induced both endocytosis and endosome recycling, whereas dynasore treatment impeded endocytosis, which subsequently impaired cell junction formation. These opposing effects were similar to those occurring in *Jmy* CKO Sertoli cells. Therefore, the loss of JMY function disorders endocytosis which in turn impairs BTB cell junction formation in Sertoli cells.

The biotinylation assay was performed to confirm that loss of JMY function does indeed disorganize endocytosis. With this procedure, membrane protein endocytosis was visualized in Sertoli cells. The results provide the first evidence that membrane proteins are continuously endocytosed and recycled back to the cell surface, resulting in an enhanced biotinylated barrier at the cell–cell interface. JMY involvement in this process was clearly documented by showing that the loss of its function interrupted endocytosis by causing the internalized protein to accumulate in the cytoplasm rather than translocate into early endosomes. Interestingly, our results indicate that JMY expression is not required for plasma membrane internalization but it is essential for endocytosed vesicle trafficking into early endosomes. Notably, the actin cytoskeleton is reported to play an important and variable role in vesicular transport (Liu, 2016; Papadopulos, 2017). This realization supports the notion that JMY induced vesicle trafficking is mediated through JMY interacting with the actin cytoskeleton.

Our protein interaction analyses provide other indications that JMY along with other actin binding proteins (i.e. Arp3, Sorbs2, α-actinin1) cooperate with each other to perform specific functions in Sertoli cells. One of them includes the Arp2/3 complex which JMY activates and produces branched F-actin filaments. Moreover, the Arp2/3 complex modulates invagination and pinching off of endosomes. They are then delivered to sorting endosomes as well as actin branches to promote BTB structural reorganization in a number of different vesicle-trafficking pathways (Lie et al., 2010). Remarkably, JMY was capable of nucleating actin filaments in either the presence or absence of the Arp2/3 complex (Zuchero et al., 2009). Herein, our results indicate that the effect of JMY on the actin cytoskeleton seems to be independent of the Arp2/3 complex in Sertoli cells. Its protein interaction might instead depend on its three WH2-domains that interact with other proteins.

To determine if this speculation is credible, IP and MS analysis were performed and revealed that JMY could interact with Sorbs2 and α-actinin1 in Sertoli cells. Sorbs2 (also known as Arg Kinase-binding Protein 2, ArgBP 2), contains an N-terminal SoHo (Sorbin homology) domain and three conserved SH3 domains in the C-terminal region. It has been reported to associate with actin fibers, adherens junction and tight junctions, and is involved in actin cytoskeletal organization, cell adhesion and migration (Roignot and Soubeyran, 2009). On the other hand, α-actinin1 (ACTN 1) is one of the major actin cross-linking proteins and anchors actin filaments to junctional structures and has an important role in some cellular biological processes such as cytokinesis, cell adhesion and cell migration (Murphy and Young, 2015). Recent evidence implicates that Sorbs2 can interact with α-actinin, subsequently binding to actin filaments (Rönty et al., 2005; Anekal et al., 2015). The current study is the first one describing interactions between the α-actinin1, Sorbs2 and JMY triad in Sertoli cells. Such interactions are consistent with our identification of colocalization of JMY with Sorbs2 and α-actinin1 along the peripheral edges of Sertoli cells. This delimited expression was particularly evident proximal to the nuclei. These regions demarcate the basal region which delineates both Sertoli cell polarity and the boundary between the uppermost apical and basal regions constitutes the BTB. Additionally, loss of *Jmy* function could lead to disordered distribution of Sorbs2 and α-actinin1, suggesting a critical role played by JMY in cross-linking the actin-related proteins to actin filaments.

In conclusion, JMY, an actin regulating protein, is essential for mediating dynamic remodeling of the BTB junctional proteins in Sertoli cells. It involvement in this process is requisite for the recycling phase of endocytic trafficking. Accordingly, JMY is crucial for sustaining the BTB structural organization commensurate with sperm health and male fertility.

## Materials and methods

### Animals

C57BL/6 mice were purchased from Shanghai Laboratory Animal Center. Anti-Mullerian hormone (*Amh*)-Cre mice (Jax number 007915) and *Jmy*-loxP mice (the *Jmy* exon 3 flanked by 2 loxP sites) were from Shanghai Research Center For Model Organisms (Shanghai, China). All of the mice were acclimated in the Animal Center of Shanghai Jiao Tong University School of Medicine. Animal experiments were conducted according to the International Guiding Principles for Biomedical Research Involving Animal, as promulgated by the Society for the Study of Reproduction. This research program was approved by the ethics committee of Shanghai Jiao Tong University School of Medicine (NO. A2015-034).

### Fertility evaluation

Individually housed, sexually mature male mice (8 weeks old) cohabitated with two virgin female mice (10 weeks old) for 7 days, and they were then separated from one another. Every day during cohabitation, females were examined for vaginal plugs as evidence of mating. Approximately, 20 days after the last day of cohabitation, the number of pups delivered by each mated female mouse was counted and the litter sizes were analyzed.

### Sperm parameters analyses

The cauda epididymides were dissected and then placed in pre-warmed (37°C) Tyrode’s Buffer (Sigma-Aldrich) to allow dispersion of spermatozoa. After 15 minutes, sperm motility, progressive motility and concentration were analyzed by computer-assisted sperm analysis (CASA) (Hamilton Thorne).

For teratozoospermia analysis, a sperm pellet was initially smeared on a glass slide. After reaching dryness at room temperature, the slide was fixed and stained as described in the Diff-Quick method (BRED Life Science Technology Inc., China). The slide was viewed under a microscope (Olympus BX53) equipped with an UPlanFN N 40×/0.75 objective (Olympus).

### Tissue preparation

Mice were euthanized with CO_2_ and then their testis and epididymis were removed. For RNA analysis, tissue was immediately snap frozen in liquid N_2_. For histological analysis, tissues were fixed overnight either in Bouin’s solution or in 2.5% glutaraldehyde buffer for electron microscopy analysis.

### Histological Analysis

Testes and epididymis fixed in Bouin’s solution were embedded in paraffin, and specimens were sliced into 5 mm thick sections and mounted on glass slides, followed by deparaffinization and rehydration. The sectioned testicular and epididymal tissues were then stained with hematoxylin and eosin (H&E) and observed under a microscope (Olympus BX53) equipped with an UPlanFN N 20×/0.5 objective and an UPlanFN N 40×/0.75 objective (Olympus).

### Transmission electron microscopy analysis

Small pieces of testicular tissue were immersed in 2.5% glutaraldehyde solubilized in 0.1M phosphate buffer (pH 7.4) for 1 day. The tissues were then fixed in 1% osmium tetroxide and dehydrated through a graded ethanol series, and embedded in Epon 618 (TAAB Laboratories Equipment). Ultra-thin sections (70-90 nm) in the seminiferous epithelium region were stained with lead citrate and uranyl acetate, and then examined at 100 kV with a Philips CM-120 (Philips).

### Western blot analysis

Mice testes or cultured cells were homogenized in RIPA lysis buffer (Thermo Fisher Scientific) containing protease inhibitor cocktail (Roche) on ice for 30 min followed by centrifugation at 12,000 xg, for 10 min, at 4°C. The proteins in the supernatant were collected and the protein concentrations were determined by the BCA Protein Assay Kit (Thermo Fisher Scientific).

Protein samples (20 μg) were separated using 8% – 16% denaturing polyacrylamide gels, then transferred to polyvinylidene difluoride (PVDF) membranes (Millipore) using a semi-dry transfer apparatus (Bio-Rad). Membranes were blocked with 5% bovine serum albumin (BSA) for 1h at room temperature and immunoblotting was performed overnight at 4°C with the primary antibodies (Table S3), followed by incubation with secondary antibody conjugated to HRP (Jackson ImmunoResearch). Signals were generated by enhanced chemiluminescence (Millipore) and detected by luminescent image analyzer (GE imagination LAS 4000).

### Immunohistochemistry (IHC) and Immunofluorescence (IF) analyses

IHC and IF stainings were performed using standard protocols. For IHC staining, paraffin sections were dewaxed and rehydrated, followed by antigen retrieval through boiling the tissue for 15 min in 10 mM citrate buffer, pH 6.0. Then, the Histostain LAB-SA Detection kits (Invitrogen) were applied according to the manufacturer’s instructions, and the sections were stained using DAB and nuclei were counterstained with hematoxylin. Digital images were captured under a microscope (Olympus BX53) equipped with an UPlanFN N 20×/0.5 objective (Olympus).

For IF staining, frozen 8-μm thick sections were prepared and then fixed with 4% paraformaldehyde for 20 min at 4°C. The unspecific binding sites were blocked with 10% BSA/PBS for 60 min at room temperature, and sections were incubated with the primary antibodies (Table S3) overnight at 4°C. Then, fluorescent-labeled secondary antibodies (1:500, donkey anti–rabbit Alexa Fluor 488, donkey anti–mouse Alexa Fluor 488, or donkey anti–rabbit Alexa Fluor 555, donkey anti–mouse Alexa Fluor 555, Jackson ImmunoResearch) were used. Nuclei were counterstained with DAPI (Sigma-Aldrich). The fluorescence signals were detected under a laser scanning confocal microscope (Carl Zeiss LSM-510, Germany) equipped with an argon laser (488 nm), a He/Ne laser (543 nm), an EC Plan-NEOFLUAR 63×/1.25 objective and a LD LCI Plan-APOCHROMAT 25×/0.8 objective (Zeiss). Digital images were captured and processed using Aim software (Zeiss Systems).

### Cell cycle analysis

Rate of germ cell development during spermatogenesis was studied in testis using propidium iodide (PI) staining combined with flow cytometer analysis, which resolved cell types based on differences in their DNA content. Briefly, the testes were incubated in 1 ml PBS containing 1 mg/ml collagenase (Type IV; Sigma) at 37 °C for 30 min with gentle agitation. After sedimentation for 5 min, the supernatant was removed and the seminiferous tubules were collected. Then, the seminiferous tubules were digested in accutase cell dissociation reagent (Innovative Cell Technologies) for 15 min and the samples were passed through a nylon filter followed by centrifugation at 400 xg for 5 min at room temperature. Thereafter, the pellet was re-suspended with 0.5 ml PBS-E (1 mM EDTA in PBS) and fixed with ice-cold 70% ethanol overnight at 4°C. The cells were permeabilized followed by staining with 0.5 mL PI/ribonuclease staining buffer (BD Biosciences) and incubated at room temperature for 15 min in the dark. Finally, DNA content of PI stained cells was analyzed in a flow cytometer (Beckman Coulter CytoFlex S) equipped with a 561nm laser. In each sample, 50,000 events were recorded and further analyzed by Cytepert software (Beckman Coulter). Three different cell populations were resolved in seminiferous tubules: 1N (spermatids), 2N (spermatogonia, secondary spermatocytes and Sertoli cells) and 4N (primary spermatocytes), and their ratios were calculated.

### BTB assay

In vivo BTB assay was performed as described (Holembowski et al., 2014). Biotin labeled reagent (10 mg/ml Sulfo-NHS-SS-Biotin, Thermo Scientific) was freshly prepared in PBS containing 1 mM CaCl_2_, and then 25 µl of this solution were carefully injected into the center of the testis at low pressure. The contralateral testis was injected with the same volume of calcium chloride solution alone and served as a negative control. After 30 min exposure to biotin, the animals were euthanized, and the testes were harvested and fixed in 10% neutral buffered formalin. Testis sections were probed sequentially with Alexa Fluor 488 conjugated Streptavidin (Jackson ImmunoResearch) to evaluate the extracellular distribution of biotin within the seminiferous epithelium. Finally, the fluorescence signal was captured under a laser scanning confocal microscope (Carl Zeiss LSM-510) equipped with an argon laser (488 nm) and a LD LCI Plan-APOCHROMAT 25×/0.8 objective (Zeiss).

### Primary Sertoli cell culture

Primary Sertoli cell culture was carried out as described (Sato et al., 2013; Holembowski et al., 2014) with some modifications. Briefly, to obtain Sertoli cells for primary culture, testes of adult mice were decapsulated in HBSS and seminiferous tubules were dispersed in a HBSS solution containing collagenase (0.1%)/hyaluronidase (0.1%)/DNase (0.04%) for 20 min at 34°C. After washing with PBS, an additional digestion step was performed with accutase cell dissociation reagent (Innovative Cell Technologies) for 15 min at 34°C. The tubular pellet was washed with PBS and Sertoli cells were freed from the seminiferous epithelium by resuspending the pellet in DMEM/F12 medium. Cells in the supernatant were collected and cultured in DMEM/F12 medium containing 5% FBS overnight at 34°C. Following attachment of the Sertoli cells to the bottom of the tissue culture plate, they acquired an irregular shape, whereas the germ cells remained suspended and were easily removed by repeated washing. The purity of primary Sertoli cells was routinely analyzed by immunofluorescent staining with SOX9.

### Endocytosis assay

Endocytosis was evaluated as described previously (Le et al., 1999). Briefly, cell surface proteins were biotinylated with 0.5 mg/ml Sulfo-NHS-SS-Biotin (Pierce) in PBS containing 0.9 mM CaCl_2_ and 0.33 mM MgCl_2_ at 4 °C for 30 min, and then quenched with 50 mM NH_4_Cl in PBS at 4 °C for 15 min and subsequently, incubated at 34 °C for the indicated periods of time during which endocytosis occurred. Biotinylated cargo proteins were then stained with Streptavidin-coupled FITC (Jackson) and detected under a laser scanning confocal microscope (Carl Zeiss LSM-510) equipped with an argon laser (488 nm) and an EC Plan-NEOFLUAR 63×/1.25 objective (Zeiss).

### Protein Immunoprecipitation (IP) and mass spectrometry (MS) assay

Protein immunoprecipitation was characterized using the Pierce Direct IP kit (Thermo scientific) according to the manual instructions. The LC-MS analysis was performed as described (Liu et al. 2015). All MS/MS spectra were searched using Proteome Discoverer2.2 software against the mouse UniProt database. Two missing cleavage sites were allowed. The tolerances of peptides and fragment ions were set at 6 ppm and 0.5 Da, respectively.

### Statistical Analysis

All data were analyzed using SAS 8.2 software, and results are presented as mean ± SD. Group comparisons were made using Student’s t-test where appropriate. One-way analysis of variance (ANOVA) test was used assuming a two-tail hypothesis with *P*< 0.05. Differences were considered statistically different when *P*<0.05.

## Supplemental material

Fig. S1 immunofluorescent staining indicates that JMY was localized in both Sertoli cells and spermatids in the seminiferous epithelium. Fig. S2 illustrates the gene engineering scheme used to obtain mice with conditional deletion of *Jmy*. Fig. S3 shows the COG analysis of proteins that immunoprecipitated with JMY. Fig. S4 shows that the testicular morphology, litter size and sperm parameters are normal in germ cell-specific deletion of *Jmy* (*Jmy* flox; *Mvh* cre) male mice. Table S1 shows results of the MS analysis of proteins immunoprecipitated with JMY. Table S2 shows the results of KEGG pathway analysis of proteins immunoprecipitated with JMY. Table S3 lists the primary antibodies used in this study.

### Acknowledgments

The authors thank Ms.Yanqin Hu (Shanghai Key Laboratory for Reproductive Medicine) for her technical assistance in histological analysis and also thank Ms. Rong Fu (Core Facility of Basic Medical Sciences) for her technical assistance in flow cytometry assay. The authors are grateful to Prof. Peter Reinach for his editorial assistance.

This research project was supported by grants from the National Natural Science Foundation of China (No. 81701503, No. 81370752 and No. 81571487) and Science and Technology Commission of Shanghai Municipality (No. 16ZR1418600).

## Author contributions

Y. Liu conducted and performed experiments, analyzed data, and prepared the manuscript. J. Fan performed the animal experiments and analyzed the data. Y. Yan, X. Dang, R. Zhao and Y. Xu performed part of molecular experiments. Z. Ding designed and supervised the project, and provided final approval of the manuscript.

## Abbreviations

ARP3: actin-related protein 3
BTB: blood–testis barrier
CKO: conditional knock out
DHT: dihydrotestosterone
EEA-1: early endosomal antigen
ES: ectoplasmic specialization
H&E: hematoxylin and eosin
IP: immunoprecipitation
MS: mass spectrometry
JMY: junction-mediating and regulatory protein
Sorbs2: Sorbin and SH3 domain containing protein 2
ZO-1: zonula occludens-1

## Tables and their legends

**Table S1. MS identification of proteins immunoprecipitated with JMY**

**Table S2. KEGG pathway analysis of proteins immunoprecipitated with JMY**

**Table S3. Primary antibodies used in this study**

**Figure S1. JMY localization in mouse testis.** JMY immunofluorescent staining was selective and evident in the seminiferous epithelium (A). The staining was greatly localized to the Sertoli cells (B) and in the post acrosomal region of spermatids in close proximity to the manchette structure (C). Arrows indicate JMY immunofluorescent staining in Sertoli cells (B) or spermatids (C). Scale Bars: (A) 50 μm; (B and C) 20 μm.

**Figure S2. Scheme of engineering *Jmy* Sertoli cell CKO mice**. Two loxP sites were inserted into both sides of the third exon in the *Jmy* gene using homologous recombination. Specific deletion of *Jmy* was carried out by a Sertoli cell specific expression of Cre recombinase.

**Figure S3. Orthologous groups of proteins clusters (COG) identified based on analysis of proteins coprecipitating with JMY in Sertoli cells.**

**Figure S4. Germ cell fertility unaltered by loss of *Jmy* function.** Male germ cell fertility is unaffected in *Jmy* CKO (A-C) Litter size (A), Sperm concentration (B) and Sperm motility (C). *Jmy* CKO male mice (*Jmy* flox; *Mvh* cre) germ cell fertility is similar to those in WT male mice. Error bars represent SD (n=5). (D) H& E stained testicular sections indicate no significant change in histological structure and integrity in *Jmy* CKO testis (n = 3). Scale Bars: (main) 100 μm; (enlarged) 50 μm.

